# Weak electric fields of deep brain stimulation can entrain spiking in multi-compartment cortical neuron models

**DOI:** 10.64898/2026.09.08.750085

**Authors:** Maud Bosman, Nina Doorn, Hil G.E. Meijer, Tjitske Heida, Bettina C. Schwab

## Abstract

**Background:** Deep brain stimulation (DBS) is widely used to treat neurological disorders, but how it exerts its therapeutic effects remains an open question. Although DBS primarily acts through stimulated subcortical structures and their networks, recent studies have demonstrated that cortical electric field (E-field) strengths generated during DBS are comparable to, and can exceed, those shown to modulate neuronal activity with transcranial alternating current stimulation. We therefore investigated whether and how these weak DBS fields can directly modulate cortical spike timing.

**Methods:** We used multi-compartment computational models of five neuron types across all cortical layers and exposed them to E-fields modeled as DBS pulses. E-field amplitudes spanned the typical range of cortical E-field strengths during DBS, while frequency and orientation were varied. Entrainment was assessed using peri-stimulus time histograms and quantified by the phase locking value (PLV).

**Results:** Weak DBS fields modulated spike timing of some neurons by either increasing or decreasing the likelihood of firing immediately following the stimulation pulse. This modulation reflected entrainment to the stimulation, with the PLV increasing with E-field amplitude and frequency. The magnitude and direction of spike-timing modulation varied across neuron types and depended on the orientation of the field.

**Conclusion:** These findings suggest that cortical E-fields of DBS may directly influence activity of some cortical neurons, alongside the established indirect cortical effects mediated by subcortical targets and their networks. This provides a new perspective on how DBS may influence cortical activity and offers insights into its potential mechanisms of therapeutic and/or side effects.

**Highlights:**

- Multi-compartment models to study how DBS E-fields affect cortical neuron dynamics.
- E-fields of DBS can directly affect cortical neuron spike timing.
- Probability of firing increases or decreases following DBS pulses.
- Some pyramidal neurons can phase-lock to weak electric fields as low as 1 V/m.
- Weak fields of DBS may play a role in therapeutic and/or side effects.

**Graphical Abstract:** 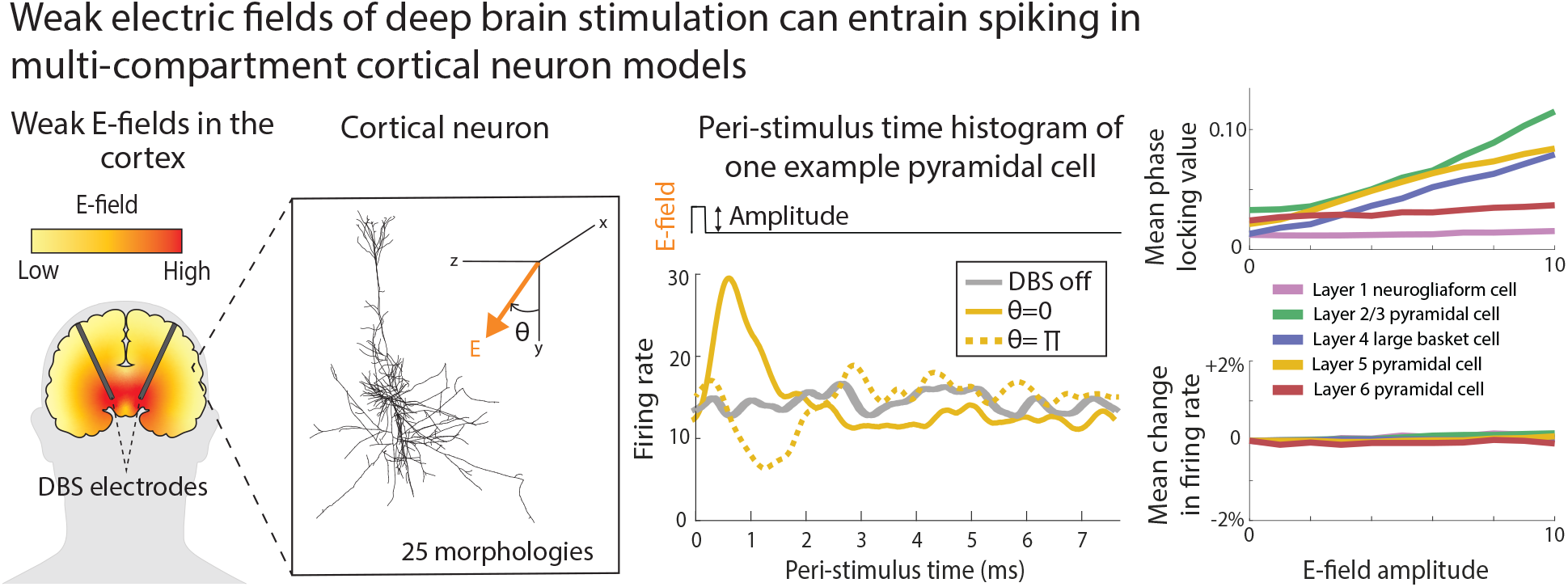

## Introduction

Deep brain stimulation (DBS) is an effective treatment for several neurological disorders, while its therapeutic outcomes and stimulation-induced side effects largely vary across patients. These effects are influenced by factors such as stimulation parameters and electrode placement [1, 2]. DBS delivers electric stimulation through implanted electrodes deep in the brain, generating electric fields (E-fields) that modulate neural activity. The effects of stimulation extend beyond the immediate target through network-mediated mechanisms, altering activity across the basal ganglia-thalamo-cortical loop [3]. Despite its widespread clinical use, the mechanisms by which DBS modulates neural activity and thereby influences clinical symptoms remain incompletely understood [4].

Current mechanistic explanations attribute the effects of DBS E-fields purely to the modulation of activity in neurons and axons close to the stimulation electrode [5]. The volume of tissue activated (VTA) is commonly used to estimate the spatial extent of this direct modulation, with a typical activation threshold of 200 V/m [6]. Within this region, DBS may exert its therapeutic effects through several mechanisms [7], including inhibition of neural somata [8, 9], disruption of information transmission within the basal ganglia [10–12], disruption of pathological firing dynamics [13–17], and antidromic activation of axonal pathways that propagate to distant brain regions [18–20]. Yet, recent E-field simulations have shown that these distant cortical regions are themselves also exposed to weak E-fields of up to several V/m [21], as schematically illustrated in Fig. 1A.

**Figure 1:**
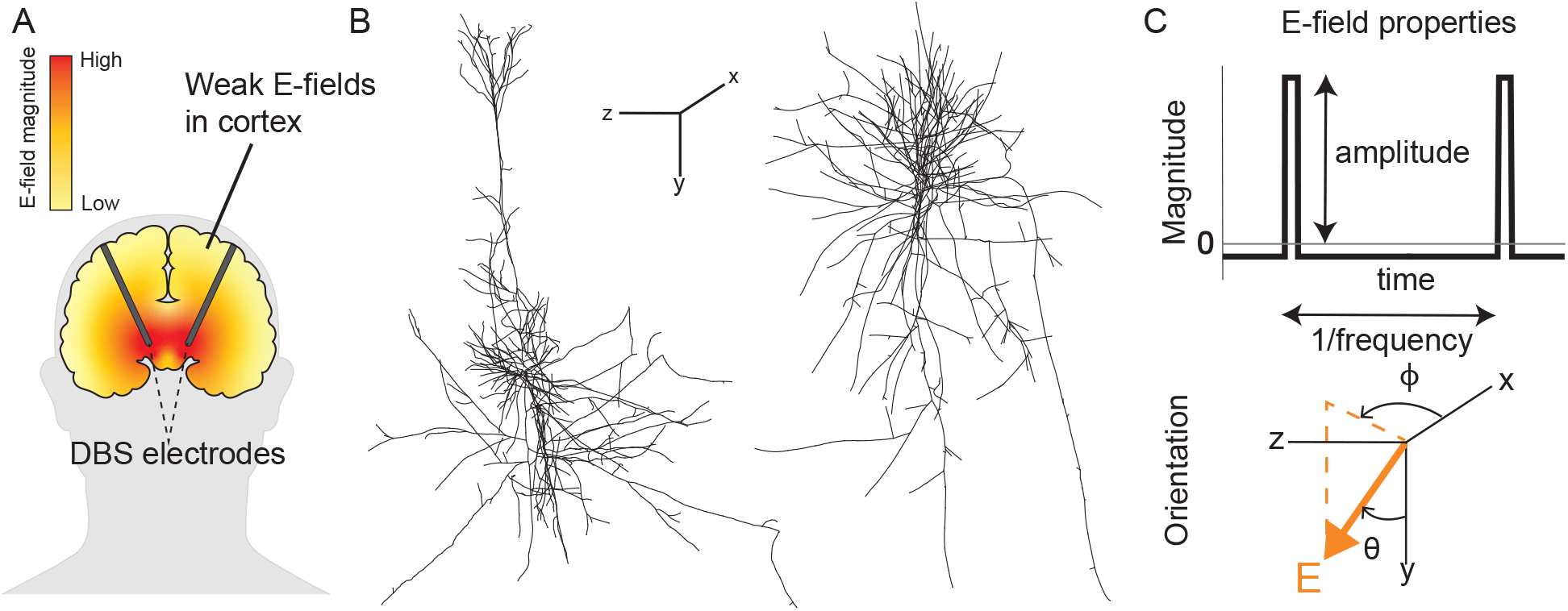
Overview of the modeling framework used to study entrainment of cortical neurons by weak DBS electric fields (E-fields). (A) Schematic illustration of weak E-fields generated in the brain during DBS. (B) Morphologically detailed multi-compartment models of representative cortical neurons [25, 27] used to simulate neuronal responses to external E-fields. Two example cells are shown out of 25 morphologies from five cell types. The orientation of the x-, y-, and z-axes is indicated. (C) Temporal waveform of the applied E-field magnitude used in the simulations, representing a DBS-like stimulus. E-field orientation relative to the neuronal morphology was defined in the x-, y-, and z-coordinate system by the azimuthal (*ϕ*) and polar angle (*θ*).

The general functional relevance of such weak E-fields has been demonstrated in studies using transcranial alternating current stimulation (tACS), which showed that weak sinusoidal E-fields can influence spike timing and neuronal entrainment without affecting overall firing rates [22–24]. Computational studies employing multi-compartment neuron models have indicated that weak E-fields can alter spike timing in a dose- and frequency-dependent manner [25, 26]. Complementary experimental studies, including single-unit recordings in non-human primates, have confirmed entrainment at E-field strengths of approximately 1 V/m [22] and 0.5 V/m [23, 24]. Importantly, E-field simulations predict that field strengths in cortical regions during DBS are comparable to, and in some regions exceed those used in tACS studies, with maximum values reaching up to 8 V/m in the orbital and insular gyrus [21].

These observations raise the possibility that weak cortical E-fields (< 10 V/m) of DBS directly modulate neuronal activity. However, because DBS delivers brief, high-frequency stimulation pulses rather than continuous sinusoidal stimulation, it remains unknown so far whether the weak cortical E-fields of DBS produce effects similar to those observed during tACS. If weak E-fields in cortical areas directly influence spike timing, they may represent an additional neural mechanism of DBS beyond the established effects on neurons and axons within the VTA and subsequent network propagation. Investigating whether and how weak DBS E-fields modulate spike timing is therefore essential to understand the full network effects of DBS.

Computational multi-compartment neuron models provide a powerful framework to investigate how E-fields modulate spike timing, as they can capture detailed neural dynamics and realistic morphologies [25, 26]. Here, we used multi-compartment neuron models to determine whether weak E-fields of DBS modulate neuronal activity and how this modulation depends on E-field parameters, such as strength, orientation and frequency. Our findings provide new insight into the potential contribution of weak E-fields outside the VTA to the mechanisms underlying therapeutic and side effects of DBS.

## Methods

### Single cell model

We employed multi-compartment, conductance-based neuron models [27], which represent neurons across all cortical layers. The original model was based on realistic neuronal morphologies reconstructed from juvenile rat somatosensory cortex [28, 29], with experimentally validated firing behaviors. Aberra et al. [27] subsequently adapted the models to reflect biophysical and geometric properties of human cortical neurons. The original data set is available on ModelDB [27] and consists of five different cortical cell types: Layer 1 neurogliaform cell (L1 NGC), Layer 2/3 pyramidal cell (L2/3 PC), Layer 4 large basket cell (L4 LBC), Layer 5 pyramidal cell (L5 PC) and Layer 6 pyramidal cell (L6 PC). Each cell type includes five morphological variants, capturing within-type variability, two examples of which are shown in Fig. 1B.

All models were implemented and simulated in the NEURON environment [30], which is designed for biophysically detailed neuron modeling. The reconstructed morphology is partitioned into many short isopotential segments. The temporal dynamics of the membrane potential in each segment are obtained by numerically integrating the cable equation:

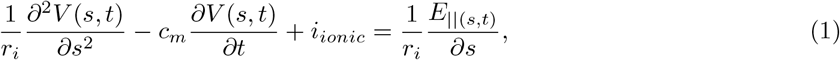

where *c*_*m*_ is the membrane capacitance, *r*_*i*_ is the membrane resistance (per unit length), *t* is time and *s* denotes the spatial coordinate along the neuronal cable. *E*_||_ is the E-field induced along the neuronal branch and *i*_*ionic*_ represents the ionic currents. The membrane and ionic current parameters were taken from Aberra et al. [27].

### Modeling of the synaptic input

To ensure baseline firing, we added a single synapse to the dendrite of each cortical neuron, following Tran et al. [25]. Synapses were placed on the basal dendrite for L1 and L4 neurons and on the apical dendrite for pyramidal neurons. We modeled presynaptic activity as a Poisson spiking train and described the synapse by a two-exponential function. We adjusted synaptic weights to yield baseline firing rates between 9.5 and 15 Hz (Table S1). This firing-rate range is consistent with reported spontaneous activity levels of cortical pyramidal neurons [31]. Standardizing baseline activity across all cell types enabled a more direct comparison of stimulation effects independent of cell-type-specific firing rates. We used a range rather than a fixed rate because exact matching was sensitive to synaptic parameter tuning, whereas small differences within this range were not expected to substantially influence entrainment to high frequencies.

### E-field modeling

We simulated the neuron’s response to an applied E-field as described by Tran et al. [25], using NEURON’s extracellular mechanism. For each compartment, we computed the extracellular potential induced by the stimulation via the quasipotential method [32]. Assuming a spatially uniform field, the extracellular potential at the *i*^*th*^ compartment is given by the inner product:

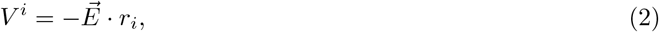

where *r*_*i*_ = (*x*_*i*_, *y*_*i*_, *z*_*i*_) denotes the three-dimensional position of the compartment and *E* is the applied E-field vector. The field orientation was specified by angles *θ* and *ϕ* such that

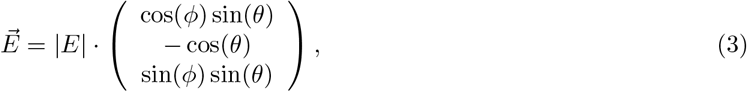

where |*E*| is the magnitude, *θ* is the polar angle measured from the negative y-axis and *ϕ* is the azimuthal angle in the x-z plane (Fig. 1C). The y-axis was aligned with the somatodendritic axis of the cells. This formulation allows the E-field to be applied along all directions in three-dimensional space. The applied E-field mimicked a typical monophasic DBS pulse (Fig. 1C) [4], with a pulse width of 240 *µ*s, selected within the range of pulse widths reported for DBS [33]. The effects of varying pulse width were additionally investigated. The pulse was followed by a low-amplitude, opposite-polarity phase of equal charge, which lasted until the onset of the next pulse. This simplified charge-balancing phase avoids assumptions about passive recharge dynamics while minimizing the contribution of the recharge phase to neuronal activation.

### E-field parameters

To investigate the effect of E-field parameters on spike timing, the E-field was varied systematically in amplitude, orientation, and frequency. First, we investigated the effect of E-field amplitude while fixing the stimulation frequency at 130 Hz and *θ* = {0, *π}*. The amplitude ranged from 0 to 10 V/m with steps of 1 V/m. Second, we examined the effect of the E-field direction by varying the polar angle *θ*, while keeping the amplitude and frequency fixed at 10 V/m and 130 Hz, respectively. The polar angle *θ* was varied from 0 to 2*π* in increments of *π/*8. Finally, we explored the effect of stimulation frequency by varying it from 30 to 170 Hz in steps of 20 Hz, for *θ* = *{*0, *π}* and an E-field amplitude of 10 V/m.

### Quantification of neuronal responses

Neuronal responses to DBS were quantified by analyzing changes in spike timing. Action potentials (spikes) were detected when the membrane potential crossed 0 mV in either a somatic or axonal compartment, with a 2.5 ms post-detection interval during which new spike detections were not registered. The detected spike times were used to compute neuronal firing rates and to quantify entrainment to the stimulation waveform. Firing rates were normalized, per neuron, to firing rates in the baseline (0 V/m) condition. To assess entrainment, we converted these spike times into their corresponding phases within the stimulation cycle. This conversion allows visualization of the phase relationship between spikes and DBS via histograms and quantification using the phase locking value (PLV). We calculated the PLV and the corresponding preferred phase (Ψ_pref_) as follows:

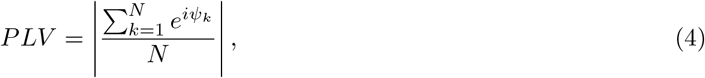

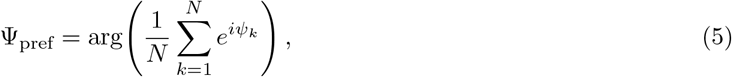

where N is the total number of spikes, *ψ*_*k*_ the phase of the k-th spike relative to the DBS waveform. The PLV ranges from zero to one, with zero indicating an absence of phase locking and one indicating complete entrainment. The statistical significance of phase locking was assessed using a permutation test. Surrogate spike trains were generated by randomly permuting the inter-spike intervals (ISIs) of the original spike train and reconstructing spike times from the shuffled intervals, thereby preserving both spike count and the ISI distribution while disrupting temporal spike alignment. A null distribution of PLVs was obtained from 10,000 iterations, and significance was assessed at *α* = 0.05.

To evaluate the average response of each cell type, spike times from all neurons within a given population were pooled and aligned to the DBS pulse. A peri-stimulus time histogram (PSTH) was constructed with 500 bins and smoothed using a Gaussian kernel with a standard deviation of 10 bin widths, using circular padding to handle edge effects. PSTHs were baseline-corrected by subtracting the baseline (0 V/m) PSTH of the same population.

To compare the wide diversity of morphologies, we used the effective length (*L*_*e*_) [25] as a simplified measure. *L*_*e*_ was calculated as the distance between the minimum and maximum coordinates along the axis of the E-field. This metric reduces complex morphologies to a comparable parameter, allowing geometric features to be correlated with simulation outcomes.

### Simulation and data analysis

Neuronal membrane potentials were simulated for two minutes, consisting of two 1-minute simulations under identical conditions with independent Poisson presynaptic spike trains. Simulations and spike detection were performed using NEURON version 8.2.6. The two 1-minute spike trains were concatenated to obtain a continuous 2-minute spike train. Time integration was performed using NEURON’s implicit (backward Euler) solver, with an integration step (*dt*) of 0.01 ms. We used a high-performance computing cluster running 8 parallel jobs (1 CPU core and 2 GB memory per job) with 5.5 hours for each simulation of 1 minute. With 2700 runs this totals 15000 CPU hours, or 1900 hours total time. We performed data analysis on a desktop computer using MATLAB R2024b.

## Results

### Weak DBS fields modulated spike timing relative to pulses

First, we tested how weak E-fields modulate spike timing of neurons at an E-field amplitude of 10 V/m, frequency of 130 Hz and the E-field oriented along the somatodendritic axis (*θ* = 0 or *θ* = *π*). These fields either increased, decreased or minimally affected the firing rate immediately after the pulse. Figure 2A-B illustrates an example neuron showing both an increase and decrease, depending on the field orientation. To characterize these effects across the population, we computed PSTHs for all individual cells and averaged them per cell-type (Fig. 2C). These were baseline (0 V/m)-subtracted to isolate stimulation-evoked firing modulation. The responses depend on the field orientation with *θ* = 0 generally associated with increases and *θ* = *π* with decreases in firing rate right after the pulse. However, responses were heterogeneous within and between cell types, with some neurons exhibiting the opposite trend or little to no modulation following stimulation.

**Figure 2:**
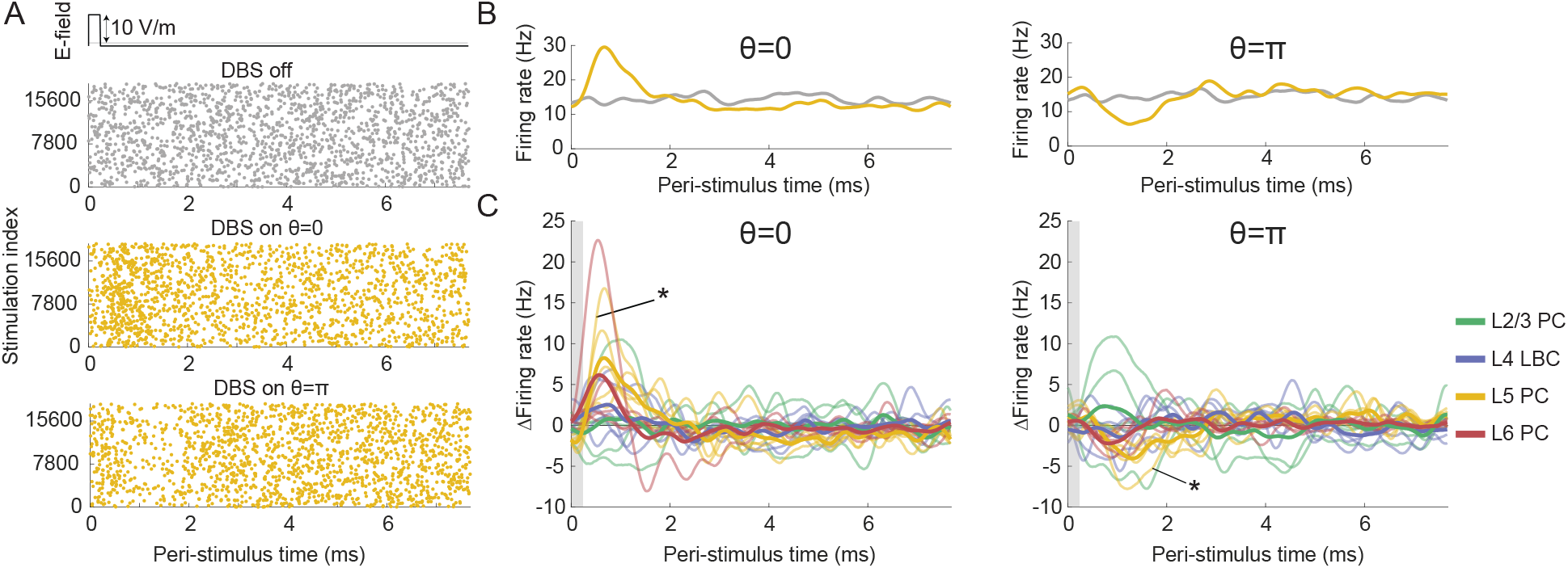
Weak DBS fields modulate spike timing relative to stimulation. Electric field was applied at 10 V/m, with a frequency of 130 Hz, and oriented along the somatodendritic axis (*θ* = 0 and *θ* = *π*) (A) Entrainment of an exemplary cell (Layer 5 pyramidal cell; cell ID 16) for two orientations (*θ* = {0, *π}*). Spike times are shown for DBS off (grey) and DBS on (10 V/m; yellow). Spike times were segmented into individual DBS cycles (1/130 Hz = 7.7 ms) and aligned to the onset of each DBS cycle, with successive stimulation cycles stacked along the y-axis (stimulation index). (B) Smoothed peri-stimulus histograms (PSTH) of both orientations (yellow) and without stimulation (grey). (C) Baseline-subtracted PSTH for individual cells (thin lines) and averaged across cell types (thick lines). *example cell shown also in figure A and B. L indicates the layer; PC, pyramidal cell; LBC large basket cell.

### E-field amplitude increased entrainment

Second, we quantified the spike timing modulation using the PLV, a measure of phase locking, and examined how this effect depended on E-field amplitude. PLV increased with the amplitude (Fig. 3A) both for *θ* = 0 and *π*. The average normalized firing rates, which reflect overall changes in the number of action potentials rather than their timing relative to the stimulus, remained stable and varied by no more than 1% (Fig. 3B). Also for individual neurons, the maximum change in firing rate was 2 % (Fig. S1). Different types of neurons exhibited distinct PLV-amplitude relationships, with high variability among neurons of the same type (Fig. 3C). Some cells displayed only minor fluctuations, whereas others showed a predominantly linear increase. A subset demonstrated an initial plateau or slight decrease in PLV before exhibiting a linear relationship. Effective length was positively correlated with PLV for both orientations, based on a Spearman rank correlation, with a significant correlation for *θ* = 0 (*r* = 0.4592, *p* = 0.0220) and a non-significant correlation for *θ* = *π* (*r* = 0.3623, *p* = 0.0758), visualized in Fig. S2.

**Figure 3:**
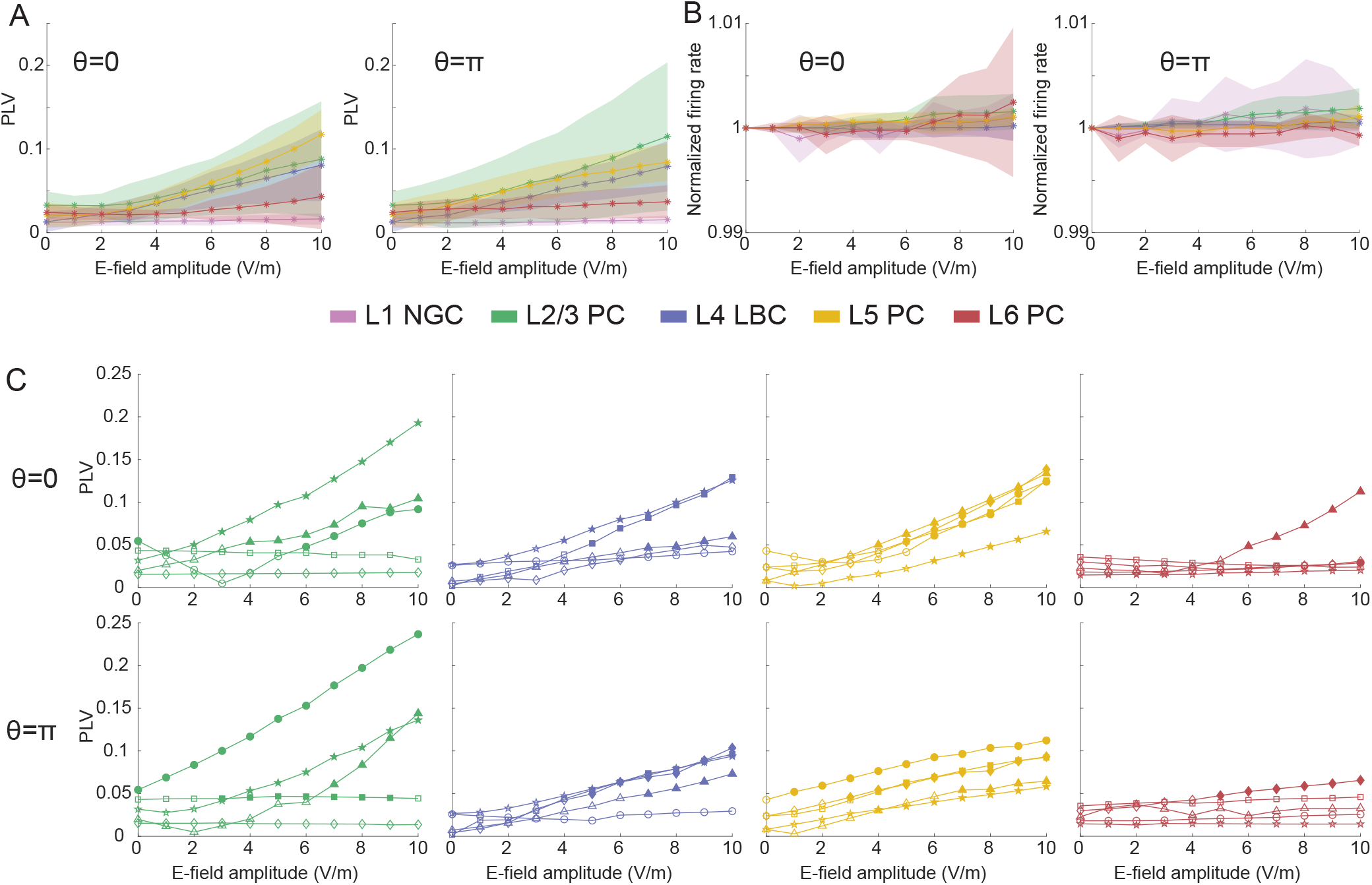
Entrainment to DBS increases with electric field (E-field) amplitude. E-field was applied at a frequency of 130 Hz, along the somatodendritic axis (*θ* = 0 and *θ* = *π*) (A) Mean phase locking value (PLV) relative to the DBS stimulation phase as a function of E-field amplitude for the five different cell types, with standard deviation indicated by shaded area. (B) Averaged normalized firing rate as a function of E-field amplitude for the five cell types, with standard deviation indicated by shaded area (C) PLV for different morphologies within cell types. Different marker shapes indicate different neuronal morphologies (five morphologies per cell type), whereas filled markers indicate significant phase locking based on a permutation test (10,000 permutations, *p <*0.05). L indicates the layer; NGC, neurogliaform cell; PC, pyramidal cell; LBC large basket cell. L1 NGC is shown in Fig. S3.

Notably, L1 NGC showed no relevant modulation by the applied E-field (Fig. S3). The phase of the pulse at which spikes occurred most consistently (preferred phase) (Fig. S4) also varied within and across neuron types and depended on E-field amplitude. Importantly, two cells exhibited a significant PLV already at 1 V/m (permutation test, *p <* 0.05) for either field orientation (*θ* = 0 or *π*). In addition, a significant PLV for either or both orientations was observed in 13 of 25 cells at an amplitude of 5 V/m and in 15 cells at 10 V/m.

### E-field orientation contributed to heterogeneous responses

Third, we examined how E-field orientation affected PLV (Fig. 4A), while keeping the amplitude fixed at 10 V/m. Given the somatodendritic alignment of the neurons along the y-axis, variations in *θ* effectively modulate the component of the E-field along this principal axis. Four neurons that did not exhibit significant phase locking at *θ* = 0 or *π* showed significant PLV at other orientations, whereas six neurons did not show significant phase locking at any *θ*.

**Figure 4:**
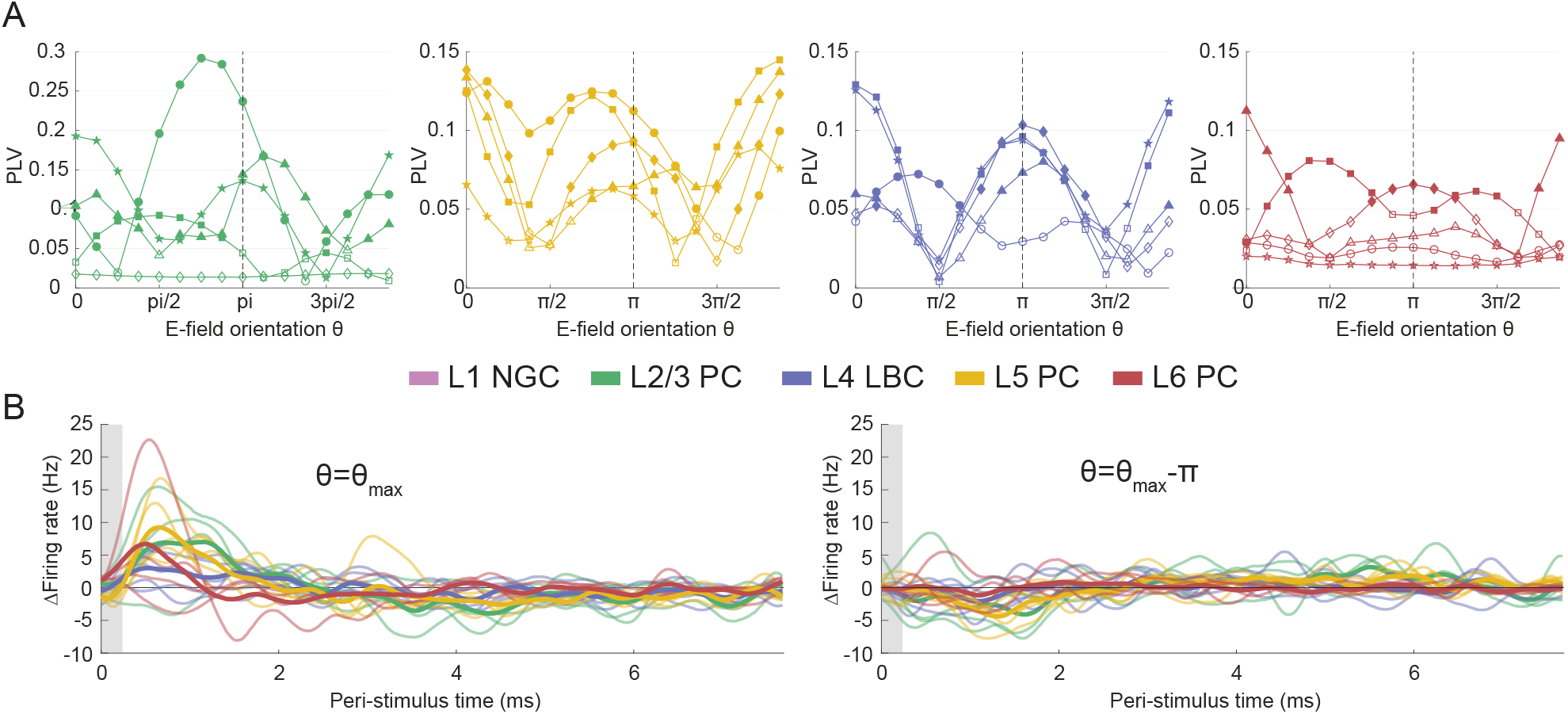
Entrainment is dependent on the orientation of the electric field (E-field) and the effective length. E-field is applied at 10 V/m with a frequency of 130 Hz. (A) Phase locking value (PLV) as a function of the E-field orientation (*θ*) for four cell types. Note that the ylimits differ between plots. (B) Baseline-subtracted peri-stimulus time histograms for individual cells (thin lines) and averaged across cell types (thick lines) for each neuron’s optimal orientation (*θ* = *θ*_*max*_) and the opposite direction (*θ* = *θ*_*max*_ − *π*). L indicates the layer; NGC, neurogliaform cell; PC, pyramidal cell; LBC large basket cell. L1 NGC is shown in Fig. S3.

### Alignment to optimal field orientation reduced response heterogeneity

Fourth, to determine whether the heterogeneity in PSTH observed for the fixed E-field orientations (Fig. 2B, *θ* = {0, *π}*) reflected differences in preferred orientation across neurons, we compared responses of these fixed and optimal orientations. From Fig. 4A, we identified for each neuron the optimal orientation that maximized significant PLV at 10 V/m (*θ*_*max*_) and its opposite direction (*θ*_*max*_ − *π*) and visualized those PSTHs in Fig. 4C. The average responses for the optimal orientations were more consistent compared to the fixed orientations, characterized by an increase in firing rate right after the pulse for *θ*_*max*_ and a decrease for *θ*_*max*_ − *π*. This effect was especially evident for L2/3 PC and L4 LBC, which displayed weaker modulation at fixed orientation compared to optimal orientations. However, variability persisted at the level of individual neurons.

### PLV increased with stimulation frequency

Finally, we investigated how stimulation frequency affected entrainment at a fixed amplitude of 10 V/m and *θ* = 0 (Fig. 5) or *θ* = *π* (Fig. S5). PLV increased with stimulation frequency (Fig. 5A) across all cell types, indicating stronger phase locking at higher DBS frequencies. For all frequencies, an increase in spiking immediately following the pulse is visible in the PSTH of L4, L5 and L6 (Fig. 5B). Notably, the amplitude of this post-pulse peak increased with frequency.

**Figure 5:**
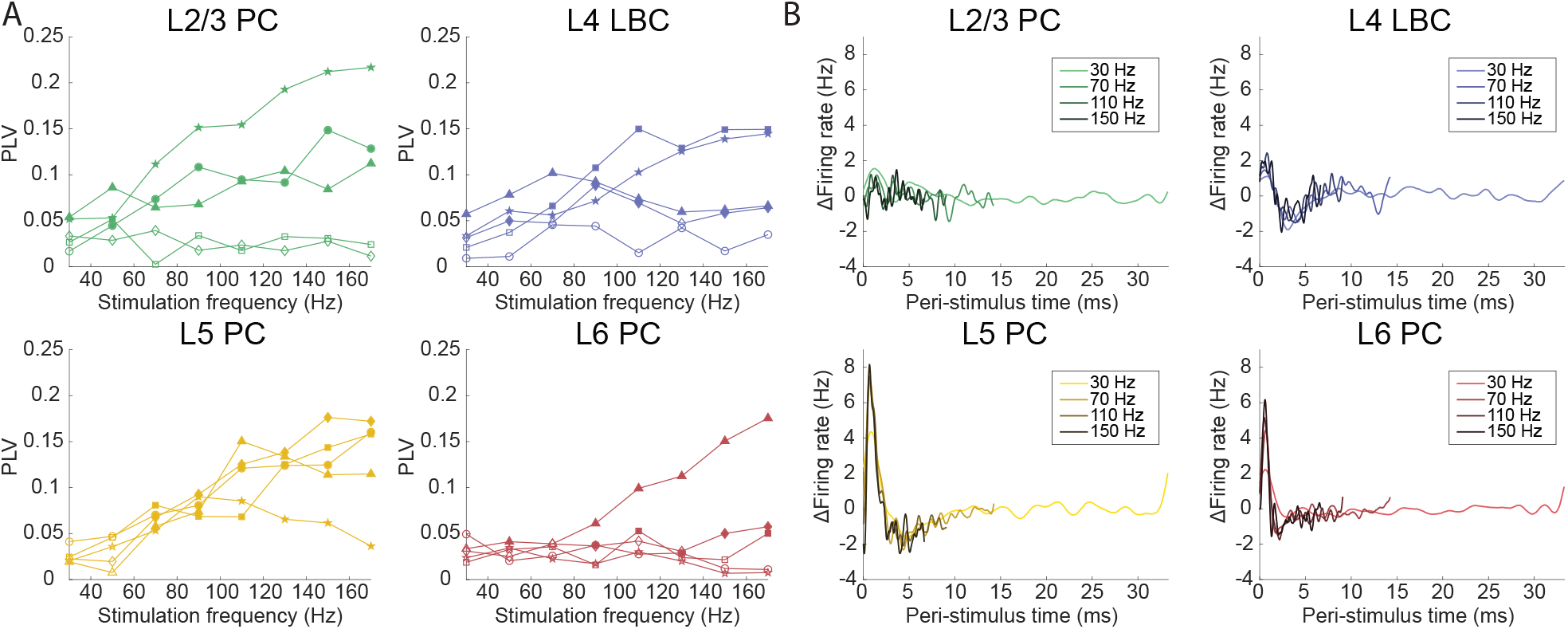
Entrainment to DBS increases with stimulation frequency. (A) Phase locking value (PLV) as a function of stimulation frequency at fixed amplitude (10 V/m) and orientation (*θ* = 0), showing an overall increase in PLV with frequency. Marker shapes indicate individual morphologies and are consistent with those used in previous figures, where filled markers indicate significant phase locking based on a permutation test (10,000 permutations, *p <*0.05). (B) Baseline-subtracted peri-stimulus time histograms for representative cell types across stimulation frequencies. L indicates the layer; PC, pyramidal cell; LBC large basket cell. Layer 1 neurogliaform cell is shown in Fig. S3. Results for *θ* = *π* are shown in Fig. S5.

## Discussion

Although the effects of DBS are generally attributed to strong E-fields near the stimulation electrode, we find that weak DBS fields (*<* 10 V/m) can also modulate neuronal activity in cortical regions. These effects are substantially weaker than the effects within the VTA and potentially also substantially weaker than indirect network effects arising from VTA stimulation, yet they may contribute to the widespread neuronal responses observed during DBS. We employed multi-compartment neuron models which showed that these weak fields can change the timing of action potentials of some cortical neurons. Importantly, this entrainment is heterogeneous, with its strength depending on the amplitude, orientation and frequency of the field. To our knowledge, this study is the first to examine how these weak DBS fields affect cortical spike timing and highlights their potential relevance to both the therapeutic and side effects of DBS.

Entrainment was already observed at our lowest tested amplitude (1 V/m), where two cells exhibited significant phase locking. While the effects of weak E-fields (*<* 10 V/m) delivered using a high-frequency (130 Hz) DBS waveform on cortical spike timing had not been directly investigated so far, studies of tACS provide relevant context. In behavioral studies, tACS has been shown to lead to strong [34, 35] and phase-specific [36] effects in some settings. Moreover, both computational and experimental studies have shown that weak sinusoidal fields (10 Hz) in the cortex, typically below 1 V/m, can entrain cortical neurons [22–26, 37]. Spike time modulation by DBS fields at the same amplitude is weaker, likely reflecting the shorter field exposure associated with DBS pulses compared with continuous sinusoidal stimulation. At the same time, cortical E-fields of DBS reach higher strengths than cortical E-fields of tACS [21].

Because orientation influences entrainment, the extent to which these effects occur in vivo depends on the spatial organization of neuronal morphologies relative to the DBS-induced E-field. Although cortical neurons are organized in columns, their orientation relative to the E-field varies across cortex. A dependence on field orientation of tACS has previously been reported in layer 5 pyramidal cells, where entrainment followed a cosine dependency on the polar angle relative to the somatodendritic axis [38]. We observed a similar relationship for DBS in a subset of neurons, including L5 pyramidal cells. Nevertheless, other pyramidal cells and large basket cells showed substantially greater heterogeneity in the orientation dependence. This can be attributed to the differences in morphology, where complex dendritic and axonal structures make it difficult to define a single effective axis of alignment. Effective length may account for part of this variability; neurons whose morphology is aligned with the E-field experience stronger polarization, leading to stronger modulation of their spiking activity [39]. Nevertheless, some neurons exhibited high effective length but low PLV, indicating that effective length alone cannot fully account for the observed heterogeneity in entrainment.

In addition to determining whether neurons become entrained, field orientation also influences how they respond. Specifically, it affects whether spike timing modulation manifests as an increase or decrease in firing rate after the pulse, which is reflected in the preferred phase of entrainment (Fig. S4). We observed a stable phase preference across amplitudes, with neurons typically locking either shortly after the pulse, corresponding to an increase in firing rate following stimulation or half a cycle later, reflecting a decrease after the pulse. The timing of this response relative to the pulse did not depend on stimulation frequency. In line with our findings, experimental tACS studies showed that neurons can lock to different phases of the stimulation cycle, but most commonly exhibit a preference around the peak or trough of the sinusoidal waveform, with phase preference that varies little across amplitudes [24, 40, 41]. These preferred phases reflect whether the applied E-field depolarizes or hyperpolarizes the neurons.

Beyond the effects observed at the level of individual neurons, our findings may have implications for network effects of DBS. Modulation by E-fields outside the VTA may act alongside, or interact with, subcortical mechanisms currently thought to mediate therapeutic effects. These E-fields may not only co-stimulate cortical areas, but also other subcortical structures, such as globus pallidus, striatum, or thalamus. Although neurons in these structures have different morphologies compared to cortical neurons, our results suggest that some subcortical neurons outside the VTA may be susceptible to entrainment by subthreshold E-fields. However, the extent of such effects in subcortical structures remains unknown, as these were not investigated here.

Nevertheless, even if only a small subset of neurons outside the VTA is affected, selective modulation of spike timing could potentially contribute to network-level effects depending on the connectivity and functional role of these neurons. For instance, weak and intermittent entrainment in cortical or subcortical regions outside the VTA, combined with more reliable entrainment in the strongly stimulated target region, may alter the relative timing of spikes along cortico-subcortical and subcortical pathways. Such shifts in spike timing relationships could, in turn, drive activity-dependent plasticity. For example, a combined DBS–transcranial magnetic stimulation (TMS) study demonstrated that the relative timing of subthalamic nucleus (STN) and primary motor cortex (M1) activation is critical for the induction of cortical plasticity, where specific STN–M1 stimulation intervals produced lasting changes in cortical excitability [42].

The ability of weak E-fields to modulate spike timing outside the VTA raises the possibility that these fields may contribute to both therapeutic effects and stimulation-induced side-effects. Interestingly, monopolar stimulation, which produces a broader E-field, induces therapeutic effects and side-effects at lower stimulation thresholds than bipolar stimulation [43, 44]. Thus, our findings raise the question of whether differences in the clinical effects of monopolar and bipolar stimulation can partly be attributed to different subthreshold E-fields, for example in cortical areas. The extent of this contribution is expected to depend on the E-field strength in the relevant cortical areas, which varies across DBS applications and disease indications due to different E-field parameters and electrode locations.

The potential relevance of these weak E-fields can be illustrated in the context of STN-DBS for Parkinson’s disease, which modulates networks involving both subcortical structures and the motor cortex [5]. E-fields during monopolar STN-DBS can reach up to 1.5 V/m in the motor cortex [21], where in our study two cells showed significant phase locking at 1 V/m. Insular and cingulate cortex are of particular interest, as they receive field strengths of up to 8 V/m, thus in the range of E-fields shown here to influence cortical spike timing of a larger fraction of neurons. The involvement of insular and cingulate cortex in cognitive processing [45] may make their unintended modulation contribute to non-motor side-effects. Previous work has reported stimulation-dependent changes in network efficiency and information transmission in these cortical regions during DBS [46, 47]. Our findings therefore raise the possibility that DBS may directly modulate neuronal activity in cortical regions such as the orbital and insular gyri through exposure to weak E-fields.

Our study has several limitations related to the neuron models that were used. First, the models used in this study did not include large pyramidal tract neurons [27], which possess long axons to subcortical targets [48]. Recent evidence from non-human primates indicates that tACS preferentially influences long-range projection pathways [49]. Consequently, the absence of these neurons in our dataset may underestimate the effect of weak E-fields on cortical cells, since they have a higher effective length. Furthermore, the E-field magnitude experienced by these axons may differ from those estimated in the motor cortex, particularly for subcortical stimulation paradigms. Second, the firing rate of the large basket cell is lower than typically observed in vivo, to facilitate comparison between cell types. However, it remains uncertain whether and how the observed responses would generalize to higher, more physiologically realistic baseline firing rates. Third, synaptic input was applied at a single location and did not change in response to the E-fields. In vivo, synaptic inputs are distributed across the neuron and may dynamically change during stimulation, which may alter the observed entrainment.

In addition to these model-related limitations, the choice of E-field parameters, particularly pulse width, may affect the observed responses. The pulse width was fixed at 240 *µs*, whereas DBS pulse widths for treatment of Parkinson’s disease are often between 60 and 90 *µ*s but can vary widely across indications, with values of up to 450 *µ*s reported for dystonia [33]. This choice was motivated by numerical considerations. With the stepsize used in the simulations (0.01 ms), shorter pulse widths would be represented by only a small number of time steps, reducing the accuracy. Nevertheless, to explore the influence of pulse width, additional simulations were performed using shorter pulse widths (at 10 V/m). These results, shown in Fig. S6, indicate similar trends but with a reduced magnitude of entrainment.

In summary, we demonstrated that some cortical neurons are sensitive to weak DBS fields, showing that even low-amplitude E-fields (*<* 10 V/m) as observed in cortical areas can modulate their spike timing while not affecting overall firing rates. Our findings particularly highlight the importance of neuronal morphology and E-field parameters in shaping entrainment effects. Future studies should embed single neurons in network models, to investigate how stimulation-induced changes propagate through neural circuits. By demonstrating that weak E-fields can directly modulate cortical spike timing, our study provides insight into how DBS may influence cortical activity beyond network effects via the VTA, with potential implications for both mechanisms of therapeutic and side effects.

## CRediT authorship contribution statement

Maud Bosman: Methodology, Software, Validation, Formal Analysis, Investigation, Data Curation, Writing – Original Draft, Visualization, Writing - Review and Editing

Nina Doorn: Supervision, Writing – Review and Editing

Hil G.E. Meijer: Supervision, Writing – Review and Editing

Tjitske Heida: Methodology, Supervision, Writing – Review and Editing

Bettina C. Schwab: Conceptualization, Resources, Data Curation, Methodology, Supervision, Writing – Review and Editing, Project Administration, Funding Acquisition

## Declaration of competing interests

The authors report no competing interests.

## Acknowledgments

This work was financed by the European Research Council (ERC StG DECODE, grant number 101116047, to B.C.S). B.C.S. further received funding from the German Research Foundation (DFG, grant number SCHW 2023/2–1) and the Dutch Research Council (NWO, grant number 22332). We would like to thank Harry Tran for his assistance in helping us understand and use his code and Thomas Keizers for valuable discussions.

## Data availability statement

Code for this study is available on Gitlab: https://gitlab.utwente.nl/bss_development/neuro/dbs-single-cell-modelling.

## Declaration of generative AI in the manuscript preparation process

During the preparation of this work the authors used ChatGPT in order to improve structure, flow and language of the manuscript. After using this tool/service, the authors reviewed and edited the content as needed and take full responsibility for the content of the published article.

## Supporting information

**Table S1:**
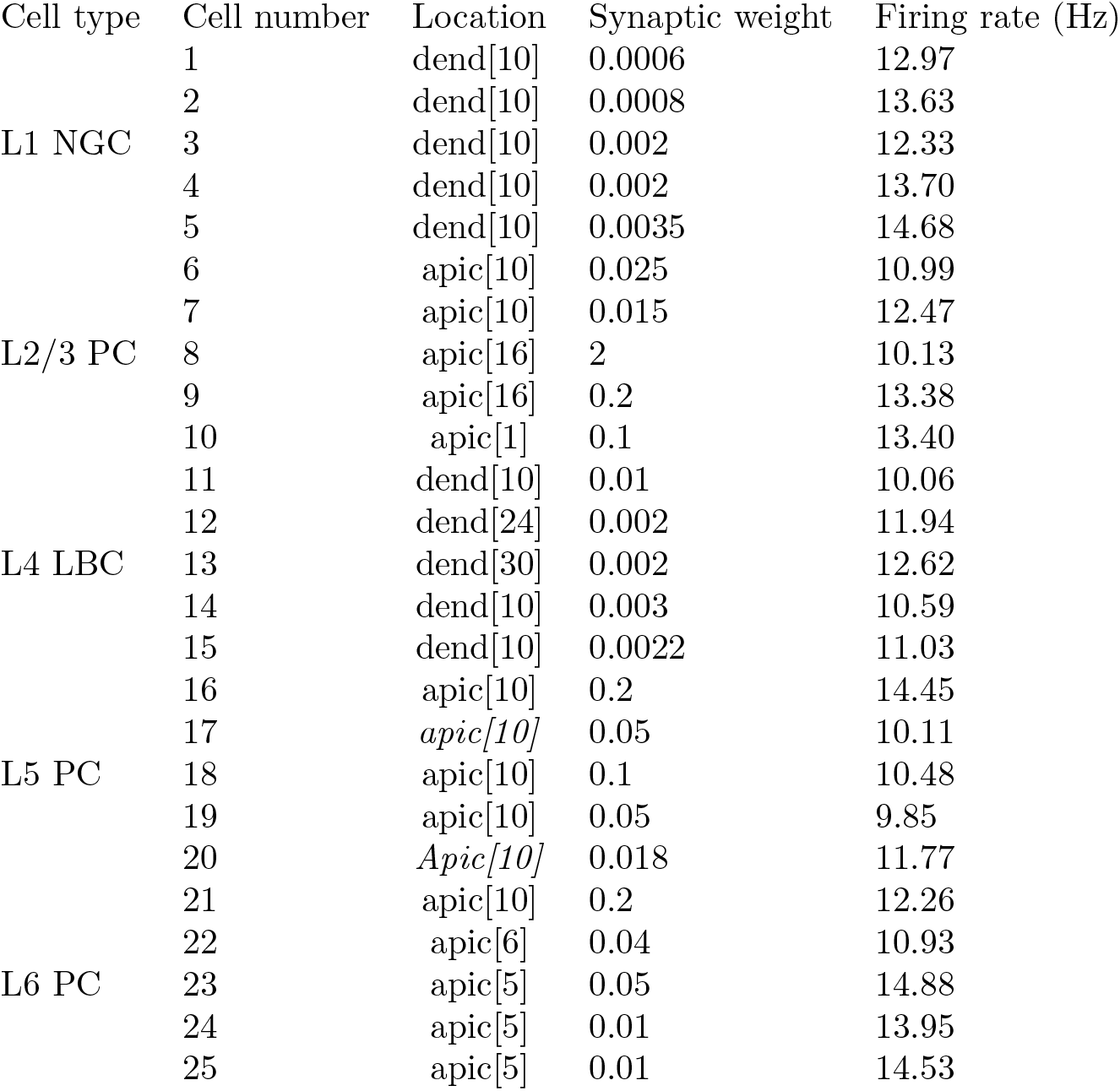
Synaptic parameter settings and resulting firing rates for each cell type. The table shows the cell type, synapse location, synaptic weight, and resulting firing rate for each cell type.

**Figure S1:**
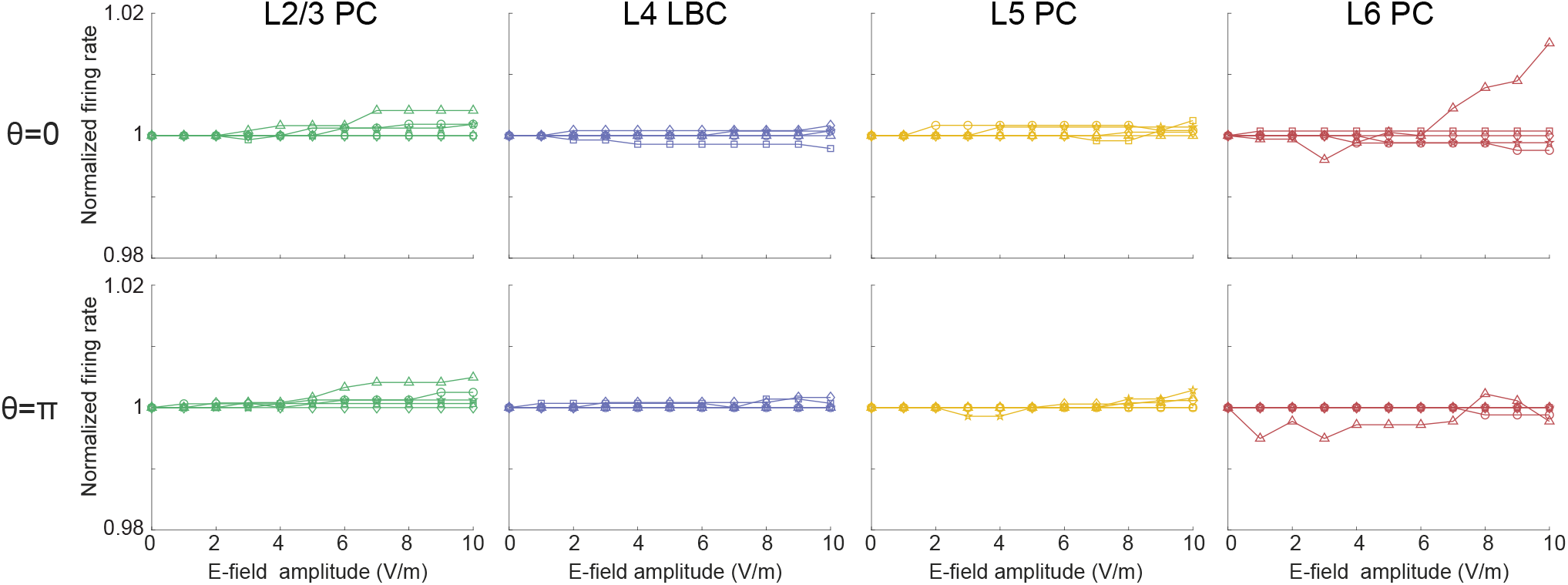
Firing rate stays constant during stimulation. Normalized firing rate as a function of E-field amplitude for different morphologies within cell types. Markers indicate different neuronal morphologies (five morphologies per cell type). L indicates the layer; NGC, neurogliaform cell; PC, pyramidal cell; LBC large basket cell.

**Figure S2:**
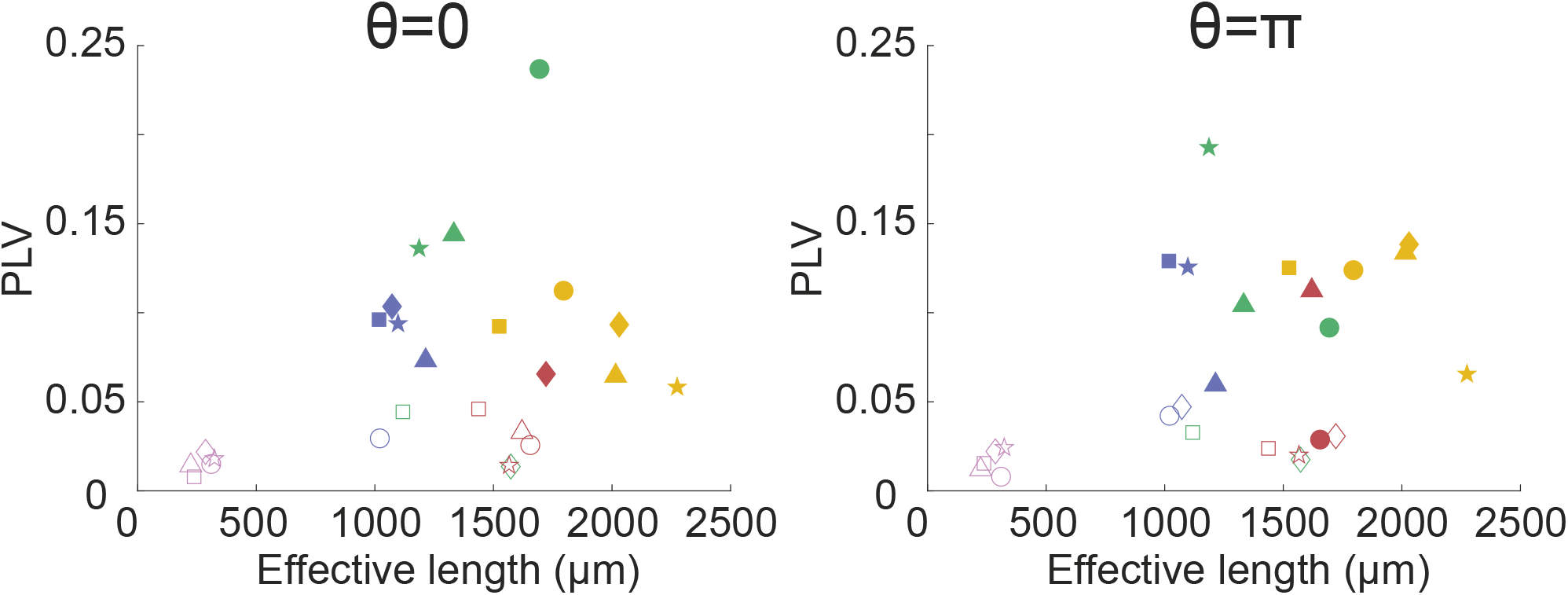
Relationship between the effective length and PLV for an E-field amplitude of 10 V/m and a stimulation frequency of 130 Hz. For *θ* = 0 there is a significant Spearman correlation (*r* = 0.4592, *p* = 0.0220), whereas for *θ* = *π* this is not the case (*r* = 0.3623, *p* = 0.0758). Marker shapes indicate individual morphologies and are consistent with those used in previous figures, where filled markers indicate significant phase locking based on a permutation test (10,000 permutations, *p <* 0.05).

**Figure S3:**
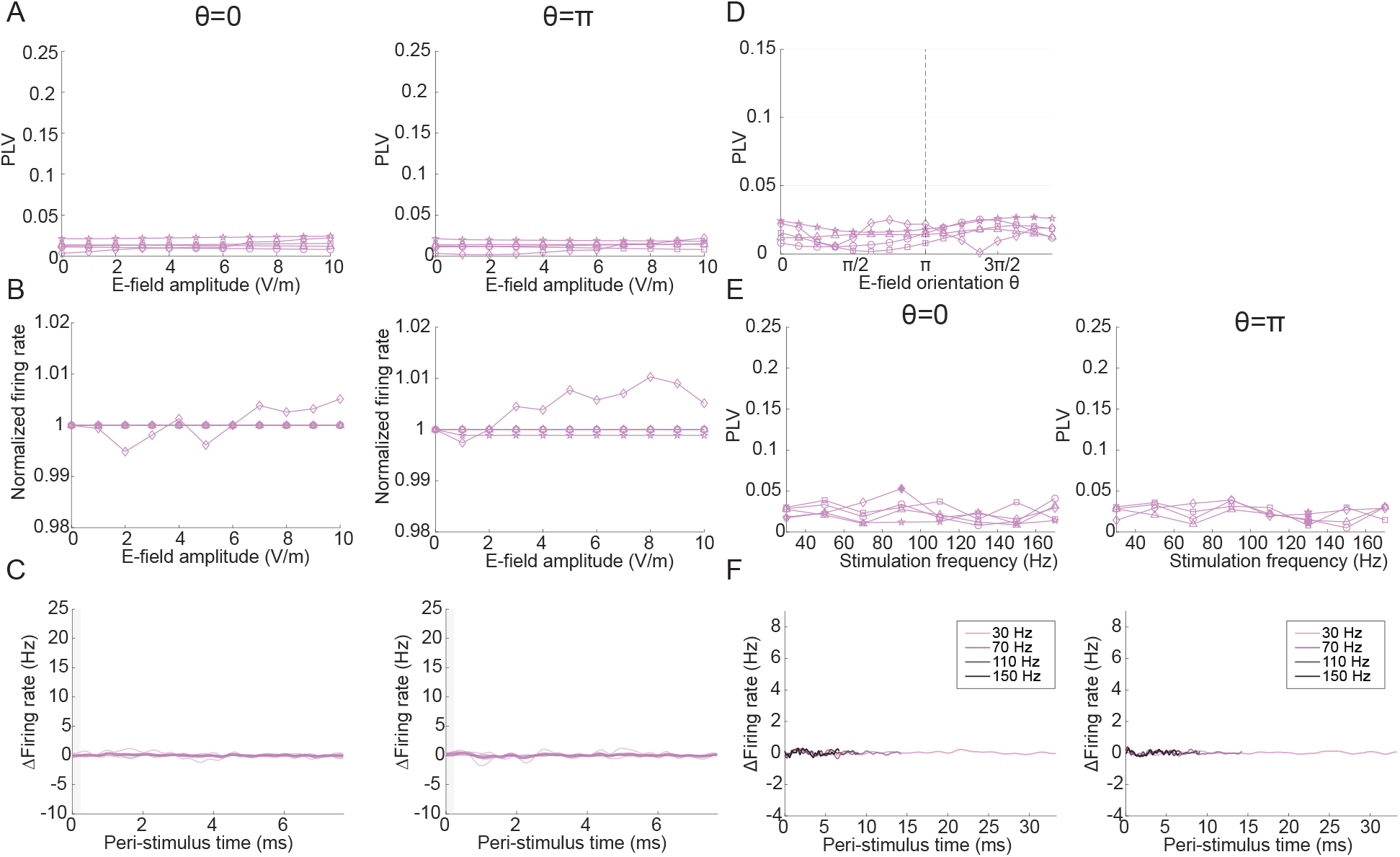
Results for Layer 1 NGC cell (L1 NGC). Markers indicate the different morphologies, and filled markers indicate significant phase locking based on a permutation test (10,000 permutations, *p <*0.05). (A-C) Electric field (E-field) was applied at a frequency of 130 Hz, along the somatodendritic axis (*θ* = 0 and *θ* = *π*), and the phase locking value (PLV) (A) and normalized firing rate (B) as a function of amplitude is shown. (C) Baseline-subtracted peri-stimulus time histograms (PSTH) for individual cells (thin lines) and averaged across cell types (thick line). (D) E-field is applied at 10 V/m with a frequency of 130 Hz. PLV as a function of the E-field orientation (*θ*). (E-F) E-field was applied at 10 V/m and fixed orientations *θ* = [0, *π*]) (E) PLV as a function of stimulation frequency (F) Baseline-subtracted PSTHs across stimulation frequencies.

**Figure S4:**
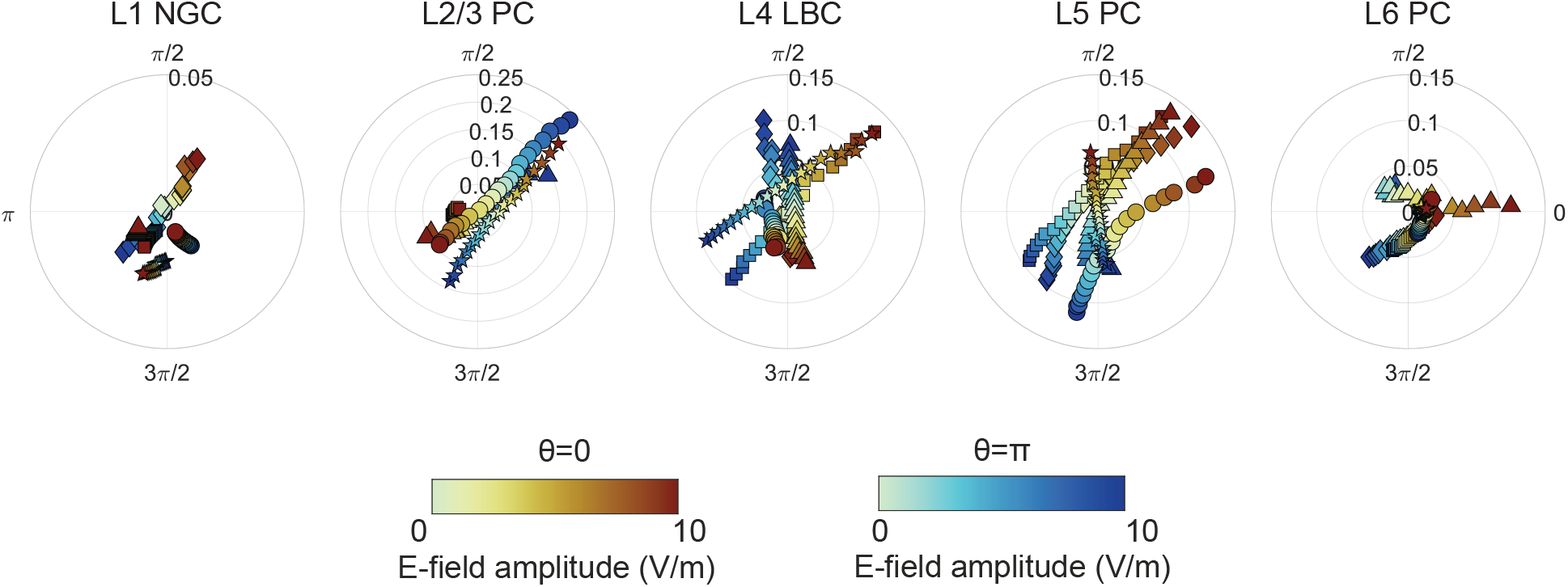
The phase that the neuron models entrain to differs across E-field amplitudes and cell types. Electric field (E-field) was applied with a frequency of 130 Hz. PLV (radius) and preferred phase (angle) as a function of E-field amplitude (marker color) for all morphologies within each cell type. Marker shapes indicate individual morphologies and are consistent with those used in previous figures. Note that the radius limits differ between plots.

**Figure S5:**
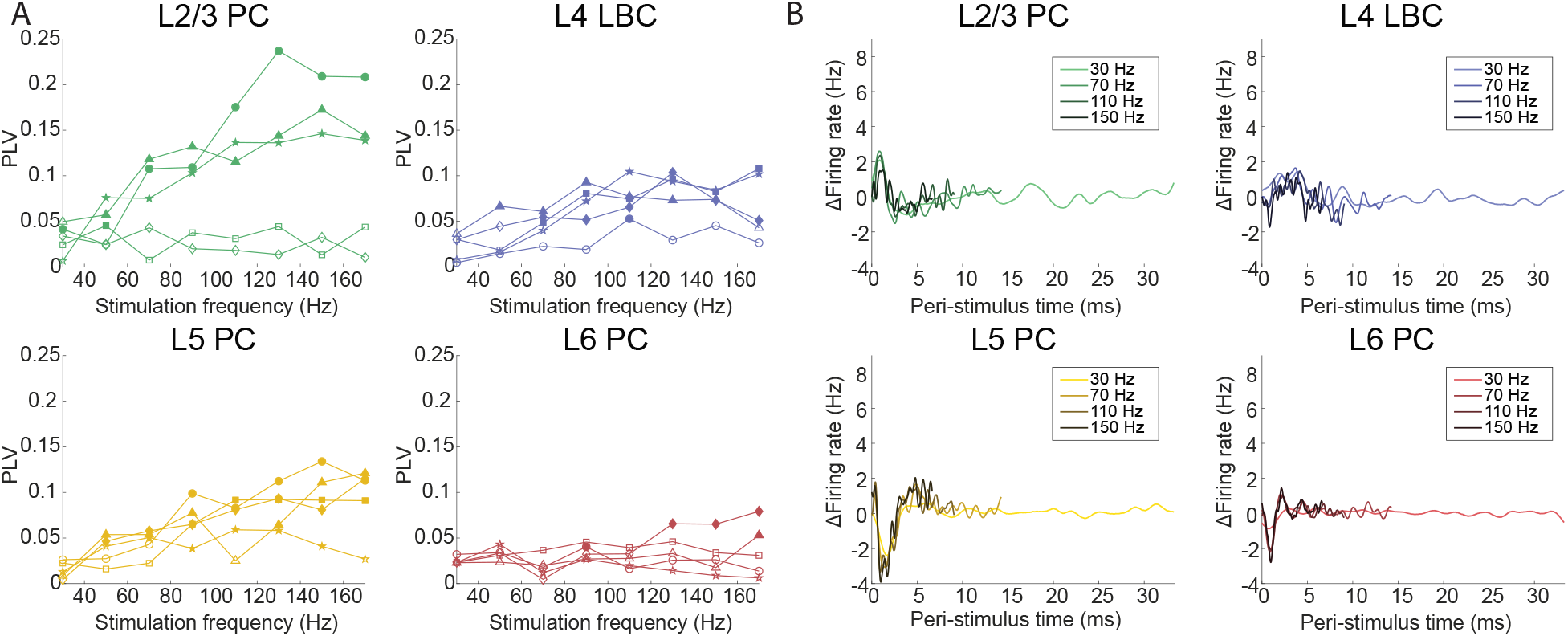
Entrainment to DBS increases with stimulation frequency, while PSTHs reveal similar responses across frequencies. (A) Phase locking value (PLV) as a function of stimulation frequency at fixed amplitude (10 V/m) and orientation (*θ* = *π*), showing an overall increase in PLV with frequency. Marker shapes indicate individual morphologies and are consistent with those used in previous figures, where filled markers indicate significant phase locking based on a permutation test (10,000 permutations, p*<*0.05). (B) Baseline-subtracted peri-stimulus time histograms (PSTHs) for representative cell types across stimulation frequencies.

**Figure S6:**
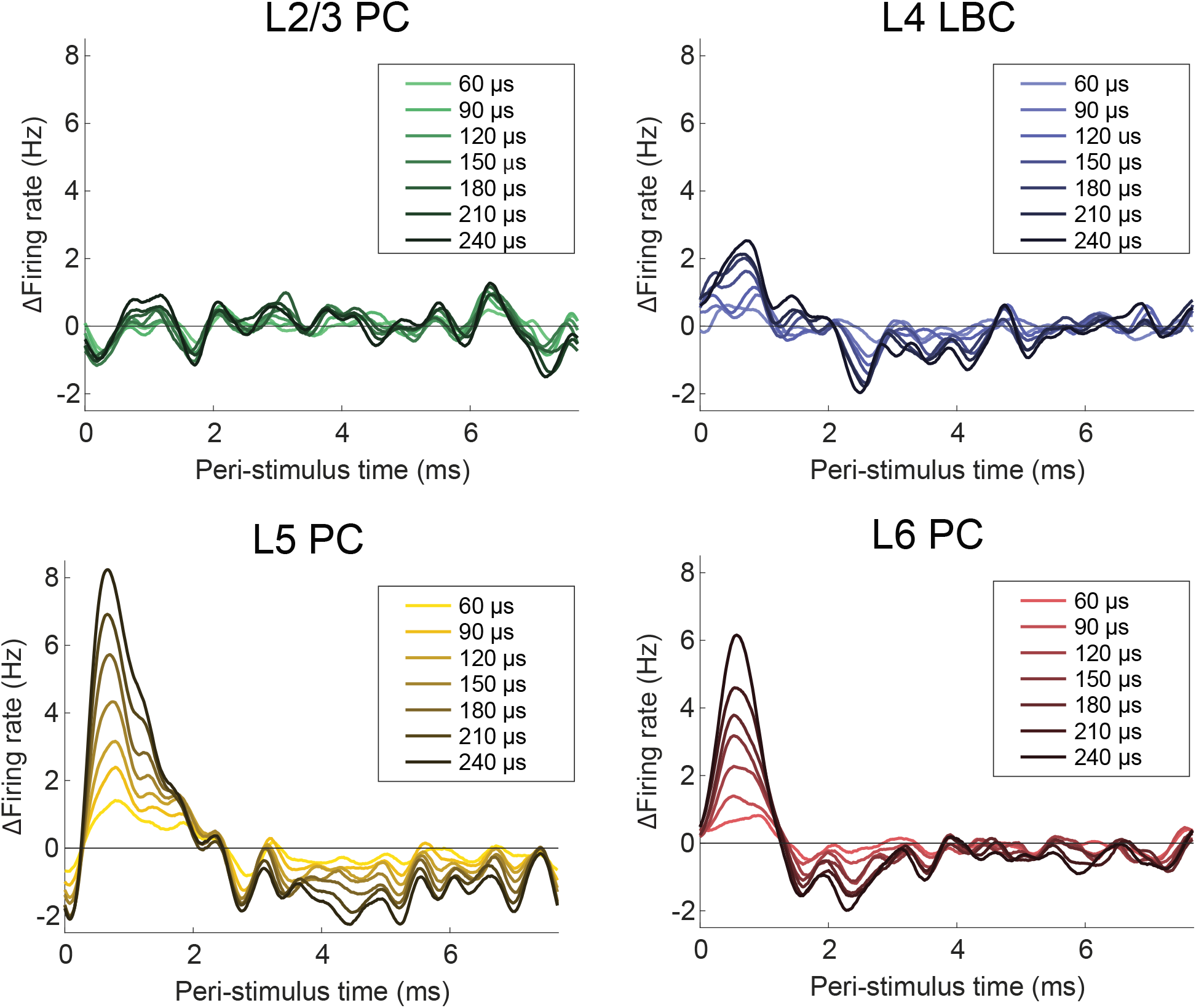
Entrainment depends on pulsewidth. Electric field was applied at 10 V/m, with a frequency of 130 Hz, and oriented along the somatodendritic axis (*θ* = 0). Baseline-substracted peri-stimulus time histograms are shown for representative cell types across stimulation pulse widths.

## Notes

https://gitlab.utwente.nl/bss_development/neuro/dbs-single-cell-modelling

### Competing Interest Statement

The authors have declared no competing interest.

